# Comparative Biochemical and Biophysical Insights into SufC ATPases from *Mycobacterium tuberculosis* and *Mycolicibacterium smegmatis mc^2^155*

**DOI:** 10.64898/2026.09.25.754312

**Authors:** Anu Ghodpage, Akhil Dinesan, Ashutosh Singh, Rajan Vyas

## Abstract

The sulfur mobilization (SUF) system is the only Fe-S cluster biosynthesis pathway in *Mycobacterium tuberculosis* (*M. tb*) and *Mycolicibacterium smegmatis* (*M. smegmatis*). In this pathway, SufC protein acts as an ATPase in the SufBC_2_D scaffold for Fe-S cluster biogenesis. This crucial role positions SufC as a promising target for the development of antimycobacterial therapeutics. However, the SufC ATPases from Mycobacterial species are not yet characterized. This study includes cloning, expression, purification, and characterization of SufC homologs from *M. tb* (*Rv1463) and M. smegmatis* (MSMEG_3124) using biochemical, biophysical, and computational approaches. Both SufC proteins were purified using affinity and size-exclusion chromatography and observed to exist as monomers in solution. Although both proteins have the conserved domains, they exhibit different ATP-dependent kinetic efficiencies, demonstrating differences in their local environment and catalytic behavior. Furthermore, CRISPR interference targeting *MSMEG_3124* results led to diminished *M. smegmatis* growth, highlighting the critical role of SufC in cellular viability. Modeling and virtual screening led to the identification of three candidate compounds, of which Z2911048515 and Z66741142 were selected for experimental evaluation. Circular dichroism spectroscopy analysis revealed slight differences in the spectral features of Rv1463 in the presence of Z2911048515 and Z66741142, while significant spectral variations were observed for MSMEG_3124. Importantly, Z66741142 was able to cause changes in the fluorescence spectra of both SufC protein homologs. Surprisingly, both compounds showed only limited inhibition of SufC ATPase activity, exhibited distinct modes of action, and may serve as preliminary chemical scaffolds for optimization and testing as effective inhibitors. This study presents functional diversity of mycobacterial SufC proteins across pathogenic and non-pathogenic species and provides a basis for developing a framework for further mechanistic and structure-guided investigation within the indispensable SUF pathway.

## 2. Introduction

Iron-sulfur (Fe-S) clusters are one of the oldest and most vital metal cofactors, consisting of iron and sulfur atoms, which occur widely throughout nature with varying architectures and functions. Some of the most commonly occurring Fe-S clusters in proteins include [2Fe-2S], [3Fe-4S], and [4Fe-4S] ^1,2^. These clusters are required for various cellular functions such as electron transport, oxidation-reduction reactions, metabolism, respiration, and DNA repair ^3–5^. In these clusters, iron ions bond with sulfide ions, forming a stable core that interacts mainly with cysteine amino acids and sometimes histidine amino acids in proteins that possess Fe-S clusters ^6^. The presence of free iron and sulfide ions is toxic for the cells, therefore, bacteria use specific multi-protein biosynthesis machinery to generate Fe-S clusters. Therefore, this machinery helps in preventing toxicity as well as the formation of these important cofactors ^7^.

In prokaryotes, mainly three Fe-S cluster assembly pathways exist: the nitrogen fixation (NIF) pathway that is necessary for the synthesis of Fe-S clusters on nitrogenase proteins, the iron-sulfur cluster (ISC) pathway that functions as the housekeeping pathway during the normal growth process, and the sulfur utilization factor (SUF) pathway, which is induced under oxidative stress, iron starvation conditions or other environmentally challenging conditions ^8–11^. Recently, phylogenomic and evolution analyses have led to the identification of two more systems for iron-sulfur cluster biosynthesis; these include the SUF-like Minimal System (SMS) in *Methanocaldococcus jannaschii* and *Methanosarcina acetivorans* (*M. acetivorans*), and the minimal iron-sulfur system (MIS) in the archaeon *M. acetivorans* and the bacterium *Helicobacter pylori* ^12^. Both SMS and MIS are considered as evolved forms of Fe-S assembly systems, demonstrating their evolutionary optimization in facilitating the assembly processes ^13^. *Mycobacterium* species, including *M. tb*, depend exclusively on the SUF pathway for the biosynthesis and maintenance of Fe-S clusters, therefore making it indispensable for survival of bacteria ^14–17^.

Comparative genomic analysis has revealed that the SUF system is highly conserved among bacteria and archaea despite the varied operon structures in various organisms. Several bacterial species, such as *Bacillus subtilis* (*B. subtilis*), *Synechococcus*, and *Escherichia coli* (*E. coli*), usually harbor a common *sufBCDS* operon, along with other accessory genes like *sufA* (an A-type Fe–S cluster carrier), *sufE*, or *sufU* (a sulfur transferase protein), and *sufT* (a DUF59-domain-containing accessory protein implicated in Fe–S cluster maturation) ^16,18–20^ (Figure 1). However, archaeal SUF operons are relatively simple and are made up of three *suf* genes, *sufB*, *sufC* and *sufD*. This implies that the SufBCD protein complex is a component of the SUF system that is evolutionarily conserved across bacterial and archaeal lineages, since it constitutes the minimum requirement for assembling Fe–S clusters under extreme environmental conditions ^21–24^.

**Figure 1.**
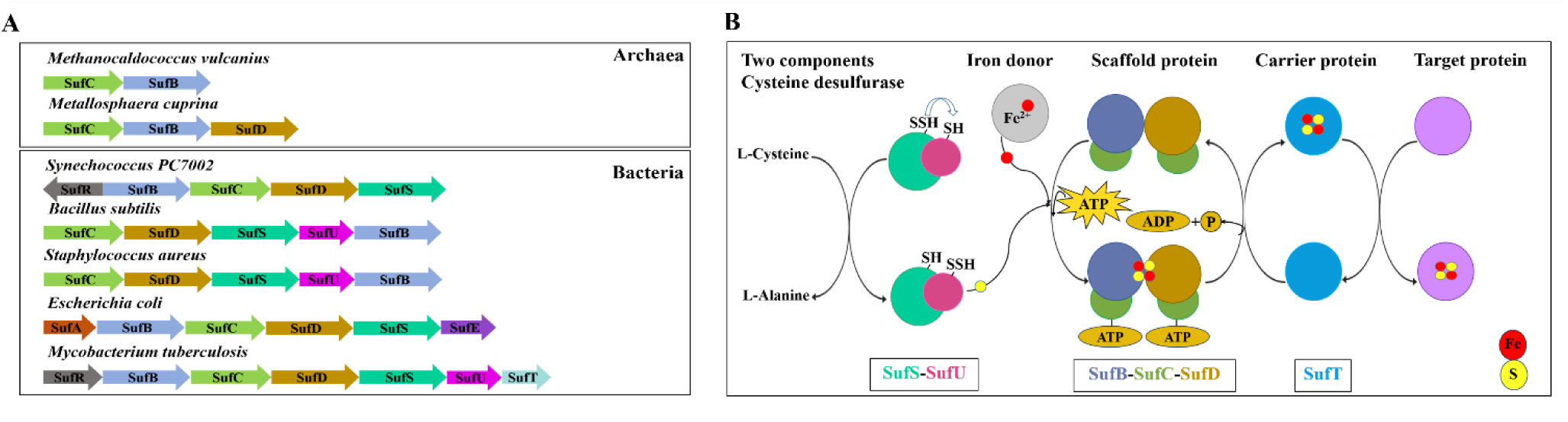
Overview of the SUF iron–sulfur cluster biogenesis system. **(A)** Comparative organization of *suf* genes across representative archaeal and bacterial species. **(B)** Schematic representation of the SUF pathway showing sulfur mobilization, Fe–S cluster assembly, and cluster delivery to target proteins.

**Table 1.** Kinetic and nucleotide-binding parameters of mycobacterial SufC proteins.

| **Protein** | **K<sub>m</sub>**<br>**(μM)** | **V<sub>max</sub> (μM**<br>**Pi/min)** | **k<sub>cat</sub> (S<sup>-1</sup>)** | **k<sub>cat</sub> / K<sub>m</sub>**<br>**(μM<sup>-1</sup> s<sup>-1</sup>)** | **K<sub>d</sub> (μM)** |
| --- | --- | --- | --- | --- | --- |
| **Rv1463-ATP** | **23.50** | **0.97** | **0.00323** | **$1.38 \times 10^{-4}$** | **22.48** |
| **MSMEG 3124-ATP** | **99.5** | **1.3** | **0.00433** | **$4.36 \times 10^{-5}$** | **3.8** |
| **Rv1463-GTP** | **165** | **0.22** | **0.000733** | **$4.44 \times 10^{-6}$** | **48.06** |

**Table 2.** Docking Score and Hydrogen-bond interactions of ATP and selected compounds.

| **Protein** | **Ligand** | **Docking Score**<br>**(kcal/mol)** | **Hydrogen-bonding residues** |
| --- | --- | --- | --- |
| Rv1463 | ATP | -7.1 | Gln94, Ser50, Gly46, Lys49,<br>Asn45, Pro44, Glu176 |
| Rv1463 | Z2911048515 | -8.3 | Gly46, Gly48, Lys49, Ser47,<br>Thr51 |
| Rv1463 | Z5027904292 | -8.4 | Gly46, Ser47, Gly48, Lys49,<br>Ser50, Thr51 |
| Rv1463 | Z66741142 | -8.3 | Lys49, Gly48, Ser50, Thr51,<br>Pro44 |
| MSMEG_3124 | ATP | -6.9 | Gly45, Gly47, Lys48, Ser49,<br>Gln93, Glu175 |
| MSMEG_3124 | Z2911048515 | -8.5 | Gly47, Ser49, Thr50 |
| MSMEG_3124 | Z5027904292 | -8.8 | Gly45, Ser46, Gly47, Lys48,<br>Ser49, Tyr61 |
| MSMEG_3124 | Z66741142 | -7.7 | Gln93, Gly45, Lys48, Ser49,<br>Tyr61 |
| 3ZDQ | ATP | -6.3 | Gly557, Arg354, Ser381 |
| 3ZDQ | Z2911048515 | -7.5 | Tyr351, Ser384 |
| 3ZDQ | Z66741142 | -7.0 | — |

In *Mycobacteria sp.,* the SUF machinery is encoded by a distinct operon comprising 7 genes, i.e., *Rv1460* to *Rv1466* in *M. tb*, and *MSMEG_3121* to *MSMEG_3127* in *M. smegmatis*. These genes encode SufR, SufB, SufD, SufC, SufS, SufU, and SufT proteins, respectively ^14^. In this machinery, the protein SufS functions as a cysteine desulfurase, removing sulfur from L-cysteine and transferring it to SufU (sulfur transferase), which subsequently participates in sulfur transfer to the SufBCD protein complex, which acts as a scaffold on which Fe-S cluster synthesis occurs. After that, SufT contributes to the maturation of Fe–S proteins, whereas SufR functions as a repressor and nitric oxide (NO) sensor, modulating operon expression in response to cellular availability of iron-sulfur (Fe-S) clusters ^14,15,20^. Interestingly, the mycobacterial operon composition differs from the classical *E. coli* SUF operon arrangement and exhibits species-specific regulatory and functional adaptations ^14,16,25^.

The SufC proteins encoded by *Rv1463* in *M. tb* and *MSMEG_3124* in *M. smegmatis* are ATPase components of the SUF system, and with ATP-binding and hydrolysis activities, are associated with the function of the SufB₂CD scaffold ^22,24,26^. Unlike classical ABC transporters, SufC is a soluble ATPase that lacks transmembrane domains but retains the conserved nucleotide binding motifs, including the Walker A and Walker B motifs, D-loop, and Q-loop, which participate in nucleotide binding, hydrolysis, and associated conformational changes ^24,26–30^. Structural analyses of SufC homologs from Gram-negative bacteria, specifically, the crystal structures of SufC from *Thermus thermophilus* (PDB ID: 2D2F) and *E. coli* (PDB ID: 2D3W), revealed a conserved ABC-ATPase fold with nucleotide-dependent structural states ^29,30^. Extending these structural studies, the crystal structures of the SufC–SufD complex (PDB ID: 2ZU0) and the SufBCD complex (PDB ID: 5AWF) elucidated the structural framework of the SufBC_2_D iron-sulfur (Fe–S) cluster assembly complex. This assembly includes two SufC proteins that interact individually with the scaffold components SufB and SufD ^23,24,31^. Furthermore, biochemical studies in *T. thermophilus* and *E. coli* have shown increased ATP hydrolysis by SufC when it interacts with SufB and SufD proteins. This interaction appears to alter the structure of its catalytic domain and, subsequently, the organization of its catalytic site ^24,29^.

The SufC protein plays a vital role in stress response and survival, as evidenced by several genetic studies across diverse bacterial species. Disruption of sufC in *Salmonella enterica serovar* Typhi resulted in decreased survival inside macrophages ^32^. Similarly, mutations affecting SufC in *Erwinia chrysanthemi* increased sensitivity to oxidative stress, disturbed iron homeostasis, and affected virulence ^26,28^. Moreover, in *Staphylococcus aureus*, antisense RNA-mediated depletion of *sufC* decreased bacterial viability, further demonstrating the importance of the SUF machinery for bacterial survival ^33^. Despite the essential role of SufC in Fe–S cluster biogenesis, its biochemical and biophysical properties in mycobacteria remain relatively poorly characterized. This study presents a comparative characterization of SufC proteins from two mycobacterial species, i.e., the pathogenic *M. tb* (Rv1463) and the non-pathogenic *M. smegmatis* (MSMEG_3124). The functional properties of SufC homologs were assessed using an integrated approach combining biochemical and biophysical characterization, CRISPRi-mediated functional analysis, and structure-guided computational analyses. Comparative analyses revealed differences in their ATP-dependent biochemical and biophysical properties, providing insights into the functional diversity of SufC proteins across mycobacterial species and contributing to a better understanding of their molecular function. These findings provide a basis for future structure-guided studies to explore SufC and the SUF pathway as potential targets for antimycobacterial drug discovery, particularly in the context of host-associated stresses that perturb Fe–S cluster homeostasis.

## 3. Materials and methods

### 3.1 Gene cloning and protein expression

To facilitate cloning, primers specific to the SufC genes were designed for both Rv1463 in *M. tuberculosis* and *MSMEG_3124* in *M. smegmatis*. In both genes, the forward primers included an NdeI restriction site, while the reverse primers contained a HindIII restriction site along with a C-terminal 6×His tag (Supplementary Table 1). For PCR amplification of *Rv1463* and *MSMEG_3124* genes, a plasmid, Rv1463-pANT7-cGST (DNASU plasmid repository), and genomic DNA extracted from *M. smegmatis mc²155* were used as template DNA, respectively. The amplified PCR products were visualized on an agarose gel, excised, purified using a gel extraction kit (Qiagen), and subjected to double digestion. Similarly, the purified pYUB1062 plasmid was digested with both restriction enzymes in a separate tube, resolved on an agarose gel, and then purified using a gel extraction kit (Qiagen). The purified, double-digested PCR product and vector were mixed with ligation buffer and T4 DNA ligase and incubated the reaction overnight at 16 °C. The next day, the recombinant plasmids were transformed into *E. coli DH5α,* and the transformed colonies obtained were screened using colony PCR and a double restriction digestion method. Positive clones were further confirmed by Sanger DNA sequencing.

To optimize protein expression, recombinant plasmids carrying Rv1463-C-His-pYUB1062 and MSMEG_3124-C-His-pYUB1062 were transformed into different *E. coli* expression strains. The expression of both proteins were optimized at temperatures ranging from 16 °C to 37 °C, with IPTG concentrations between 0.1 mM and 1 mM and post-induction incubation periods between 5 and 16 h. Based on the optimization results, *E. coli* C43 (DE3) and Rosetta (DE3) were selected for recombinant expression of Rv1463 and MSMEG_3124 proteins, respectively. For primary culture, a single colony was inoculated into Luria-Bertani (LB) broth medium containing Hygromycin (100 μg/ml) and grown at 37 °C overnight. For secondary culture, 1 L of LB medium was inoculated with 1% of the primary culture and incubated at 37 °C until the OD_600_ reached 0.6–0.8. The cells were then induced with 0.5 mM IPTG and incubated at 30 °C for 5 h. The induced cultures were harvested by centrifugation at 10,000 ×g for 10 min and stored at −80 °C till further use.

### 3.2 Protein purification

For the purification of Rv1463 protein, the cell pellet was resuspended in the lysis buffer containing 20 mM Tris, pH 8.0, 200 mM NaCl, 5 mM imidazole, 5 mM BME, 1 mM PMSF, 5% glycerol (Buffer-A). Cell suspension was kept on ice and lysed by sonication (Qsonica) at 30% amplitude with a 10 s on/off pulse for 30 min, and the lysates were clarified by centrifugation (10,000 rpm, 30 min, 4 °C). The supernatant was loaded onto Ni-NTA resin, washed with Buffer-A supplemented with imidazole at 5-30 mM, and eluted with elution buffer containing 20 mM Tris, pH 8.0, 200 mM NaCl, 5 mM BME, 300 mM imidazole, 5% glycerol (Buffer-B). For the purification of the MSMEG_3124 protein, most steps were similar to those for Rv1463, except that the lysis buffer (Buffer A) contained 10 mM imidazole. After affinity purification, the purity of both proteins was assessed by SDS-PAGE. For further purification, the eluted proteins were separated by size-exclusion chromatography using a HiLoad 16/600 Superdex 200 pg, equilibrated with 20 mM Tris, pH 8.0, 100 mM NaCl, and 2% glycerol (Buffer-C). The protein eluted from each peak was analyzed by SDS-PAGE to assess purity. To assess the effect of ATP on oligomeric state, the size-exclusion chromatography was performed with and without ATP. Both proteins, Rv1463 and MSMEG_3124, at a concentration of 1 mg/ml, were incubated with 1 mM ATP in the presence of MgCl₂ for 30 min and then loaded onto the Superdex 75 Increase 10/300 GL column equilibrated with Buffer-C. The elution profiles obtained in the presence and absence of ATP were compared to assess the oligomeric state of both proteins.

### 3.3 Circular Dichroism (CD) Spectroscopy

Purified Rv1463 and MSMEG_3124 proteins were buffer-exchanged into 10 mM sodium phosphate, pH 7.4, using a PD-10 (Cytiva Lifesciences) desalting column before analysis. CD spectra were recorded on a JASCO J-1500 spectropolarimeter at 25 °C in a 2 mm path-length quartz cuvette at a final protein concentration of ∼10 µM. Far-UV spectra were collected from 200–260 nm with a scan speed of 50 nm/min and a data interval of 1 nm. Each CD spectrum represents the average of independent measurements, with each scan obtained by accumulating three instrument scans. The blank was measured using the buffer recorded under identical conditions and subtracted from the sample spectra. All measurements were performed in triplicate using independently prepared protein samples to assess reproducibility. The secondary structure content was estimated from the far-UV CD spectra using the BeStSel web server. To examine ligand-induced changes in secondary-structure elements, far-UV CD spectra were recorded for the Rv1463 and MSMEG_3124 proteins in the presence of ATP alone or the compounds selected after virtual screening, Z66741142 and Z2911048515. For ATP-bound and compound-bound protein experiments, samples were supplemented with 25 µM ATP and 25 µM compounds and incubated for 10 minutes before measurement. The thermal unfolding measurements were also performed using the same buffer conditions and protein concentration as those used for Far-UV CD measurements. Ellipticity at 216 nm and 222 nm was monitored over the temperature range of 5-95 °C at a controlled heating rate of 5 °C/min.

### 3.4 Intrinsic Tryptophan Fluorescence Spectroscopy

Intrinsic fluorescence measurements were performed to monitor conformational changes and nucleotide- and compound-induced effects. Purified Rv1463 and MSMEG_3124 proteins were buffer-exchanged using a PD-10 (Cytiva Lifesciences) desalting column into 10 mM sodium phosphate, pH 7.4 buffer, and diluted to a final concentration of ∼5 µM. Samples were excited at 280 nm, and emission spectra were recorded from 300 – 450 nm using a 5 nm excitation/emission slit width on a JASCO J-1500 spectropolarimeter at 25°C. Measurements were recorded for apo protein, followed by stepwise titration of ATP and compound from 0-25 µM, with samples equilibrated for 10 minutes before each measurement. The fluorescence emission maximum (λmax) and the changes in fluorescence intensity were analyzed for comparison. All experiments were performed in triplicate, with independent measurements.

### 3.5 ATPase activity assay

ATPase activity of Rv1463 and MSMEG_3124 was quantified using a malachite green–based phosphate detection assay (Sigma-Aldrich, USA) according to the manufacturer’s instructions. Reactions were performed in 20 mM Tris (pH 8), 100 mM NaCl, and 5 mM MgCl₂, with varying ATP concentrations (0–1000 μM) and SufC concentration of 5 μM, as reported earlier ^26^. Reactions were incubated at 37 °C for 40 minutes, terminated with malachite green reagent, and absorbance was measured at 660 nm. Kinetic parameters Km, Vmax, and kcat were determined by fitting the initial velocities to the Michaelis–Menten equation ^34^. We also conducted enzymatic in presence of two compounds identified through virtual screening. Both compounds were solubilized in 100% DMSO and subjected to serial dilution, with ATP hydrolysis estimated across a concentration range of 0 to 1000 μM.

### 3.6 Microscale Thermophoresis (MST) assay

The binding affinities of ATP and GTP toward Rv1463 and MSMEG_3124 were determined by microscale thermophoresis (MST) using a Monolith NT.115 instrument (NanoTemper Technologies, München, Germany). Both Rv1463 and MSMEG_3124 proteins were fluorescently labeled using the RED-tris-NTA His-tag labeling dye (NanoTemper Technologies). For labeling, 200 nM protein was mixed with the labeling dye and incubated for 30 min at 25 °C. For nucleotide-binding measurements, 16-point twofold serial dilution of ATP or GTP was prepared starting from highest concentration of 1 mM. Each nucleotide dilution was mixed at a 1:1 volume ratio with 200 nM labeled protein, resulting in a final protein concentration of 100 nM in each capillary. The mixtures were incubated for 10 min at 25 °C and subsequently loaded into glass capillaries. Thermophoresis measurements were performed at 25 °C using an MST power of 20%. Changes in normalized fluorescence (F_norm) were recorded across the nucleotide concentration series, and binding curves were generated by plotting the change in normalized fluorescence (ΔF_norm) against the corresponding nucleotide concentration. Equilibrium dissociation constants (Kd) were determined by fitting the binding curves using the software provided with the MST instrument (MO.Affinity Analysis software, NanoTemper Technologies). ATP and GTP-binding measurements were performed independently for both Rv1463 and MSMEG_3124.

### 3.7 CRISPRi-mediated silencing of MSMEG_3124

To investigate the functional importance of *MSMEG_3124* in *M. smegmatis*, CRISPR interference (CRISPRi)-mediated gene silencing was performed using the pLJR962 plasmid. Three single-guide RNAs (sgRNAs) targeting *MSMEG_3124* were designed, with predicted fold-repression values of 216.7, 145.2, and 82.2 respectively ^35^. The sgRNA with the highest predicted fold repression (MSMEG_3124_N1; 216.7) was selected and cloned into the pLJR962 plasmid for subsequent gene-silencing experiments. The pLJR962 plasmid without the *MSMEG_3124*-targeting sgRNA was used as the control. Since pLJR962 carries a kanamycin-resistance marker, kanamycin (50 µg/mL) was used for selection and maintenance of plasmid-containing cells.

For the growth assay, *M. smegmatis* strains carrying either the *MSMEG_3124*-targeting CRISPRi construct or the pLJR962 control plasmid were cultured in the presence of 50 µg/mL kanamycin and grown to an OD₆₀₀ of approximately 0.6. The cultures were then serially diluted from 10⁰ to 10⁻³, and 5 µL of each dilution was spotted onto agar plates containing increasing concentrations of tetracycline (25, 50, 100, 200, and 500 ng/mL) ^36^. Plates without tetracycline were included as uninduced controls. Following spotting, the plates were incubated at 37 °C for 3–4 days ^37^. The effect of CRISPRi-mediated *MSMEG_3124* silencing on bacterial growth was evaluated by comparing the growth between *MSMEG_3124*-targeting CRISPRi strain and pLJR962 control strain across the tetracycline concentration gradient.

### 3.8 Structural Modeling

Three-dimensional structural models of Rv1463 and MSMEG_3124 were generated by homology modeling using the SWISS-MODEL server ^38^. The quality of the predicted models was evaluated using MolProbity and QMEANDisCo scoring metrics ^39,40^. To evaluate the Rv1463 model, a structural superposition was performed with the SufC homolog from *T. thermophilus* (PDB ID: 2D2F) determined by X-ray crystallography.

### 3.9 Molecular Docking and Virtual Screening

A focused ligand library comprising ATP, ADP, and small drug-like molecules retrieved from the PubChem and Enamine databases was prepared for docking analysis. Protein structures were prepared by removing the solvent molecules, adding polar hydrogen atoms, and converting them to PDBQT format using PyRx ^41^. Molecular docking was performed using AutoDock Vina as implemented in PyRx. The docking grid was centered on the ATP-binding cleft, identified through structural superposition with PDB ID: 2D2F, using a grid box of 18 × 18 × 18 Å. ATP and ADP were initially docked to validate the docking setup and identify the nucleotide-binding pocket. Initial virtual screening of the compound libraries was performed with an exhaustiveness value of 8. The top-ranked compounds were subsequently re-docked using exhaustiveness values of 16, 32, and 128 to evaluate the reproducibility and consistency of the predicted binding poses. Docked poses were visually inspected using PyMOL (The PyMOL Molecular Graphics System, Schrödinger, LLC) and Biovia Discovery Studio Visualizer 2020 (BIOVIA, Dassault Systèmes, Waltham, MA, USA) to confirm correct placement within the nucleotide-binding pocket and to assess key protein–ligand interactions ^42,43^.

### 3.10 Molecular Dynamics Simulation

Molecular Dynamics (MD) simulations were performed using Desmond software to investigate the stability and dynamic behavior of the protein-ligand complex for the control Rv1463 with ATP and Rv1463 bound to drug molecules ^44^. All systems for MD simulations were solvated in a cubic box using the TIP3P water model with a 10 Å buffer. Ions were strategically placed to neutralize the system, and the OPLS-2005 force field was used to parameterize it. Simulations were first performed for the 10 drug molecules selected through virtual screening, with ATP as a control, for 50 ns, saving the trajectory every 1.2 ps. Then, the best 3 ligands were selected and subjected to MD simulations for 500 ns, along with ATP as a control, MSMEG_3124, and the human analog (3ZDQ). Simulations were conducted under NPT ensemble conditions at 300 K and 1 atm pressure using the Nose-Hoover thermostat and Martyna-Tobias-Klein barostat. The default relaxation protocol of Desmond was used to equilibrate the system before running the MD simulation. The trajectory analysis was performed using Desmond’s simulation interaction diagram, which calculates various properties and measurements for MD simulations. Structure stability was evaluated using root mean square deviation (RMSD), while residue-level flexibility was calculated using root mean square fluctuations (RMSF). Protein-ligand interactions, such as hydrogen bonds, hydrophobic interactions, ionic interactions, and water bridges, were measured throughout the simulations to assess stability and identify which interactions persisted throughout the simulations, while the radius of gyration (Rg) and solvent-accessible surface area (SASA) were analyzed to identify ligand exposure to the solvent ^45^.

## 4. Results

### 4.1 Cloning, expression, and purification of SufC proteins

The genes encoding *M. tb* SufC (Rv1463) and its *M. smegmatis* homolog (MSMEG_3124) were amplified and cloned into pYUB1062 expression vector with a C-terminal 6×His tag. Both recombinant plasmids were verified by restriction digestion and subsequently confirmed by Sanger sequencing. For protein expression, *E. coli* C43 (DE3) and *E. coli* Rosetta (DE3) strains were used for Rv1463 and MSMEG_3124, respectively. Both proteins were first purified by Ni–NTA affinity chromatography. To enhance purity, the protein purified by affinity chromatography was subsequently loaded onto the size-exclusion chromatography (SEC). The proteins eluted by both affinity and SEC showed a predominant band at approximately 30 kDa on SDS-PAGE, consistent with the anticipated molecular weight of Rv1463 and MSMEG_3124 (Figure 2). The SEC elution profile confirmed that both proteins exist as monomers in solution. In bacteria, the two SufC monomers bind to SufB and SufD proteins, constituting a critical complex known as SufBC_2_D, which is essential for the biosynthesis of iron-sulfur (Fe-S) clusters. However, the mechanism by which SufC interacts within this complex—specifically whether it undergoes self-dimerization or relies on interactions with SufB and SufD—remains to be elucidated. To further explore the dimerization dynamics of SufC proteins, both Rv1463 and MSMEG_3124 proteins were incubated with ATP and Mg²⁺, then loaded onto the analytical SEC column. The SEC results showed no alterations in the elution profiles of Rv1463 and MSMEG_3124, indicating they remained monomeric in solution. These results indicate that ATP and Mg²⁺ do not induce detectable self-dimerization of isolated Rv1463 or MSMEG_3124 (Supplementary Figure 3A & 3B).

**Figure 2.**
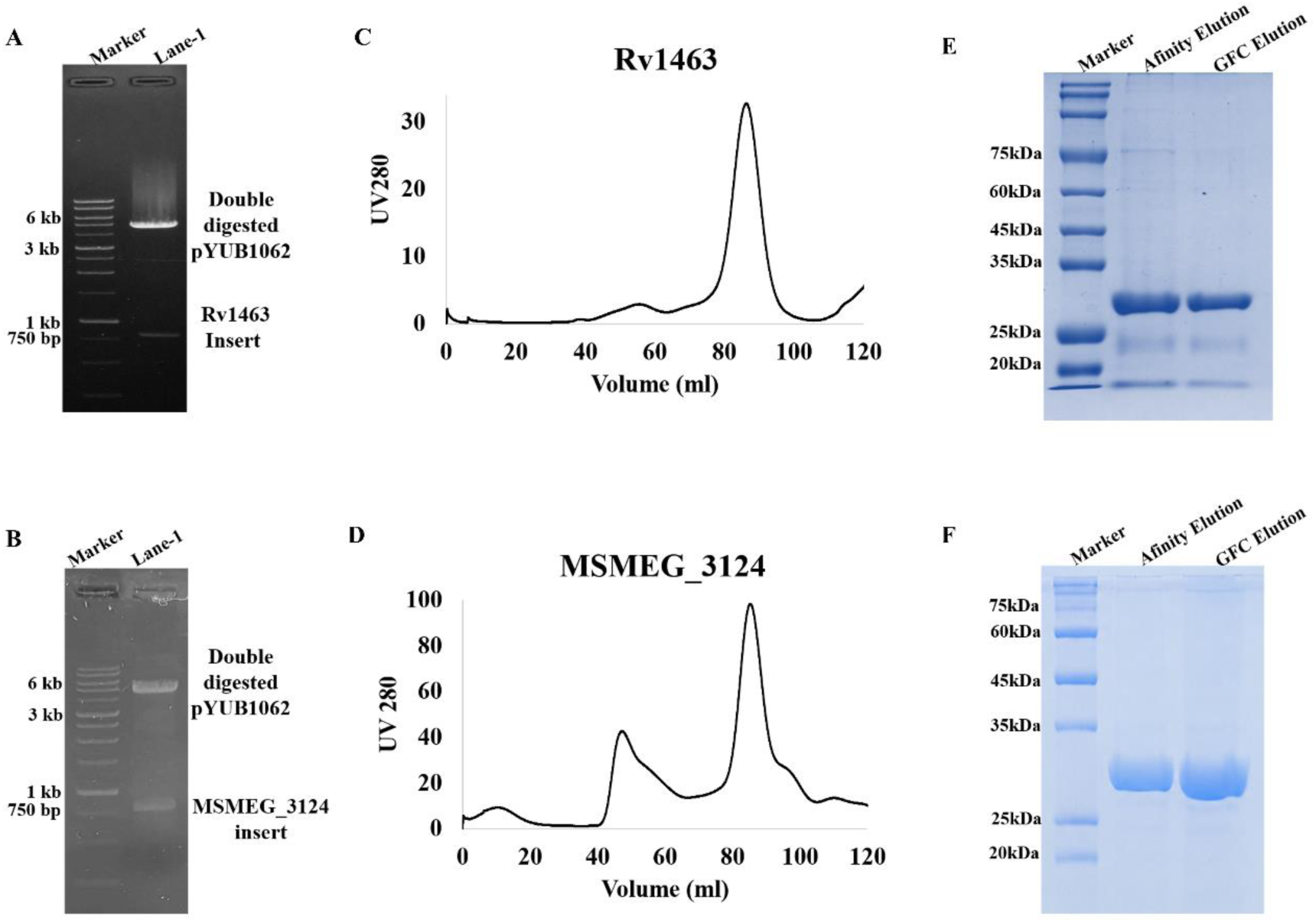
Expression and purification of mycobacterial SufC homologs. **The agarose gel shows the c**onfirmation of the recombinant plasmids carrying **Rv1463** (A) and **MSMEG_3124** (B) by restriction digestion analysis. The gel filtration chromatography profiles of purified Rv1463 (C) and MSMEG_3124 (D) proteins. After purification through Ni²⁺-affinity chromatography and subsequent size-exclusion chromatography (SEC), the purity of the proteins Rv1463 (E) and MSMEG_3124 (F) was assessed via SDS-PAGE analysis.

### 4.2 Circular Dichroism (CD) Spectroscopy

The secondary structural components of Rv1463 and MSMEG_3124 were investigated using far-UV circular dichroism (CD) spectroscopy. Both proteins exhibited pronounced negative ellipticity minima at 208 and 222 nm, suggesting that Rv1463 and MSMEG_3124 are composed of approximately 35% and 70% α-helical content respectively (Figure 3A). Moreover, thermal denaturation was evaluated by monitoring the ellipticity at 222 nm over the temperature range of 5–95 °C. Both proteins exhibited temperature-dependent changes in ellipticity, and nonlinear fitting of the thermal denaturation profiles yielded apparent melting temperatures (Tm) of 63.32 °C for Rv1463 and 57.11 °C for MSMEG_3124. The higher Tm observed for Rv1463 than for MSMEG_3124 indicates greater thermal stability of Rv1463 (Figure 3B & 3C).

**Figure 3.**
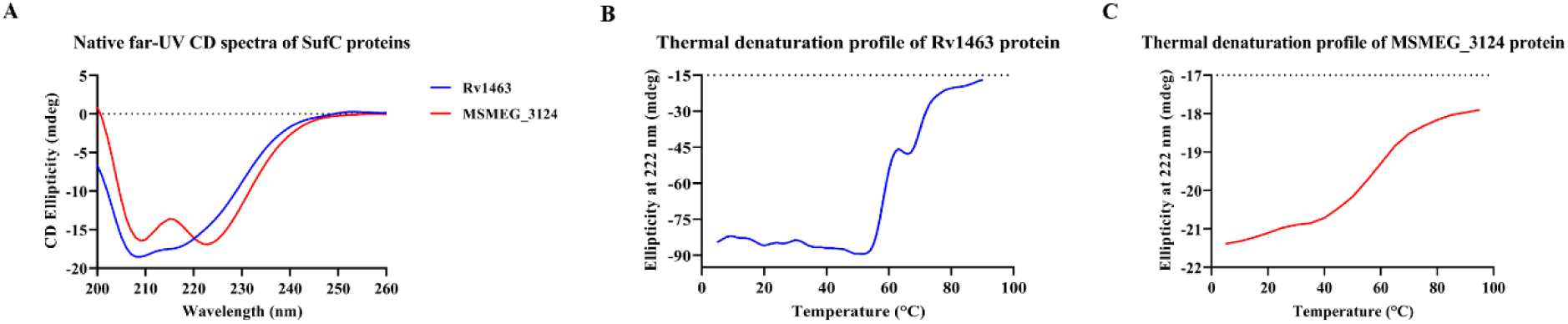
Circular dichroism and thermal denaturation analysis of mycobacterial SufC proteins. Far-UV CD spectra of purified Rv1463 and MSMEG_3124 (A). The thermal denaturation profiles of Rv1463 (B) and MSMEG_3124 (C), observed due to changes in CD ellipticity at 222 nm with increasing temperature.

To investigate the ligand-induced modifications in the secondary-structure characteristics of the Rv1463 protein, CD spectra were measured under various conditions: in the presence of ATP alone, with the compounds Z66741142 and Z2911048515 individually, and in combination with ATP. However, no significant alterations in secondary-structure content were detected across these conditions. Surprisingly, MSMEG_3124 exhibited significantly distinct CD spectra after incubation with either compound (Figure 10C & 10D). In the presence of any of the compounds, the α-helical secondary structure content of MSMEG_3124 decreased from 70 to 30%, and the addition of ATP to the above solution further resulted in a slight change in the CD spectra (Figure 10).

### 4.3 Intrinsic Tryptophan Fluorescence Spectroscopy

Intrinsic tryptophan fluorescence spectroscopy was used to examine ligand-induced changes in the local environment of aromatic residues in Rv1463 and MSMEG_3124. Titration of both proteins with increasing concentrations of ATP resulted in only minor changes in the fluorescence spectra. Similarly, the addition of Z2911048515 produced minimal changes in the fluorescence profiles of both SufC homologs. In contrast, Z66741142 induced pronounced concentration-dependent alterations in the fluorescence spectra of both Rv1463 and MSMEG_3124, characterized by a decrease in fluorescence intensity accompanied by changes in the spectral profile (Figure 11). These observations indicate that Z66741142 induces substantial changes in the local environment of tryptophan residues in both SufC homologs.

### 4.4 ATPase activity of *M. tb* SufC and its homolog

The ATPase activity of the SufC homologs was evaluated using a colorimetric assay that detects the release of inorganic phosphate (Pi) upon nucleotide hydrolysis. Besides ATP, other nucleotides including GTP, CTP, and ADP were used to assess the kinetics of Rv1463 and MSMEG_3124 enzymes. The results indicated that in the presence of ATP, Rv1463 showed an apparent Km of approximately 23.5 µM and a Vmax of approximately 0.97 µM Pi/min. In contrast, substituting ATP with GTP led to a marked decrease in enzymatic activity, with an apparent Km of roughly 164.5 µM and a Vmax of about 0.22 µM Pi/min. The other two substrates, CTP and ADP, showed only minimal phosphate release and thus are not used to determine the reliable kinetic parameters.

However, the ATP-dependent activity of MSMEG_3124 yielded a Km of ∼99.5 µM, and a Vmax of ∼1.3 (μM Pi/min). Surprisingly, compared with Rv1463, MSMEG_3124 exhibited a higher apparent Km and Vmax values. Rv1463 displayed an approximately fourfold lower apparent Km for ATP, whereas MSMEG_3124 exhibited an approximately 1.3-fold higher Vmax. Moreover, the enzyme efficiency index (Vmax/Km) indicates that Rv1463 converts substrate faster at low concentration than MSMEG_3124. For both proteins, ATP supported the highest ATPase activity among the substrates used. However, GTP showed lower activity than ATP, while CTP and ADP exhibited minimal activity (Figure 4). The ATPase assay was also performed with two compounds, Z66741142 and Z2911048515, selected through virtual screening, to evaluate their effect on the ATPase activity of both proteins. The assays were performed in the presence of increasing concentrations of Z66741142 and Z2911048515 dissolved in DMSO. Both compounds produced an approximately 20% reduction in ATPase activity, indicating partial rather than complete inhibition of ATP hydrolysis in Rv1463 and MSMEG_3124 proteins (Supplementary Figure 5).

**Figure 4.**
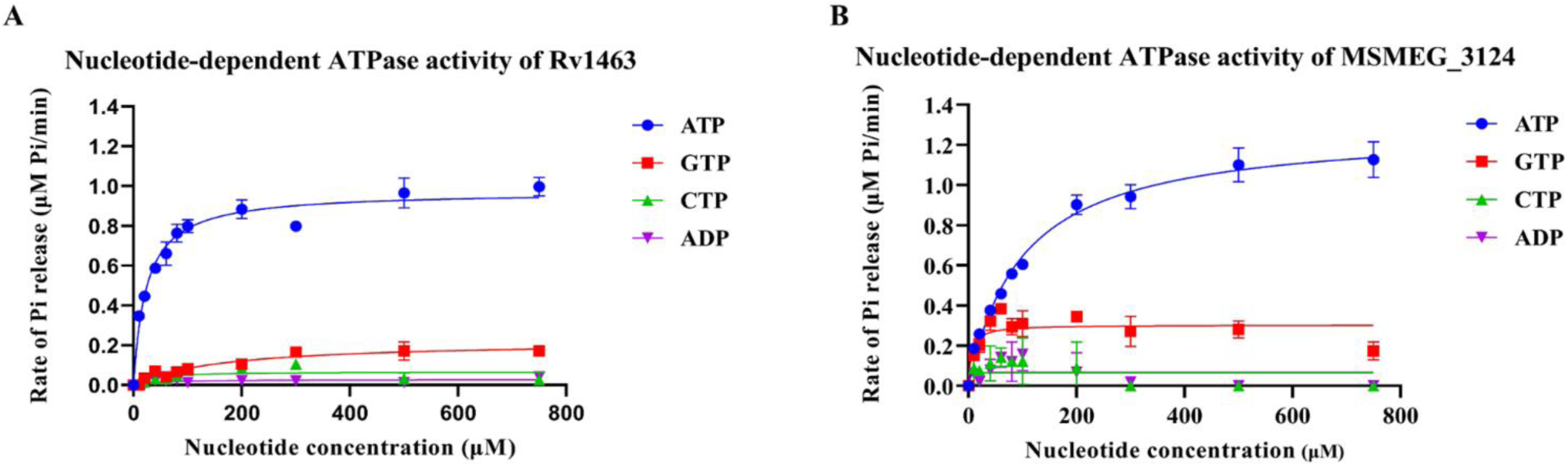
Kinetic analysis and nucleotide specificity of mycobacterial SufC ATPases. **(A)** Nucleotide-dependent ATPase activity of Rv1463 in the presence of ATP, GTP, CTP, and ADP. **(B)** Nucleotide-dependent ATPase activity of MSMEG_3124 in the presence of ATP, GTP, CTP, and ADP. The kinetic parameters were obtained by fitting the data to the Michaelis–Menten equation. Data are presented as the mean ± SD from three independent experiments.

### 4.5 Microscale Thermophoresis (MST) assay

To determine the binding affinity of SufC homologs for ATP and GTP, MST was performed. Rv1463 bound ATP with an apparent K_d_ of ∼23.5 µM, whereas GTP exhibited a lower binding affinity with an apparent K_d_ of ∼50 µM (Figure 5, Supplementary Figure 2). Both nucleotides generated concentration-dependent binding curves that were fitted using a one-site binding model. Similarly, MSMEG_3124 bound ATP with an apparent Kd of ∼ 4 µM, whereas GTP exhibited a lower binding affinity with an apparent Kd of ∼277 µM.

**Figure 5.**
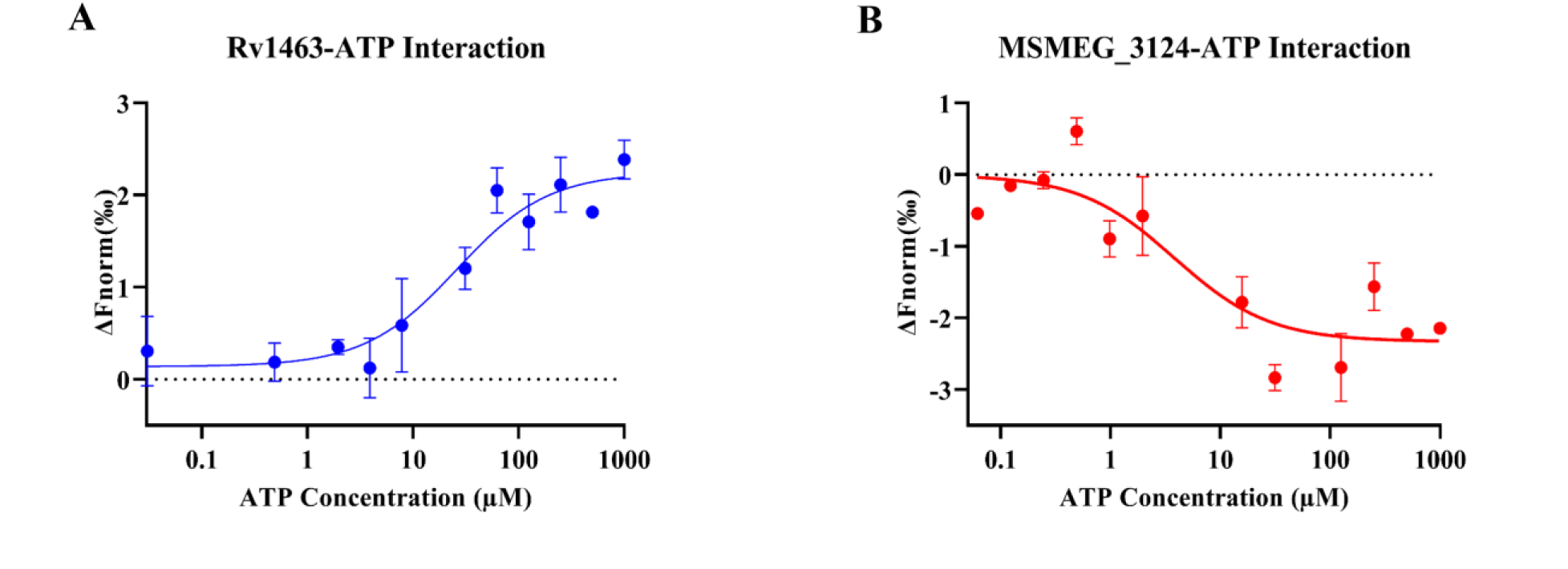
ATP binding by SufC homologs determined by microscale thermophoresis (MST). The MST profile of ligand ATP with Rv1463 (A) and MSMEG_3124 (B). The change in normalized fluorescence (ΔFnorm) was plotted as a function of nucleotide concentration, and Kd values were obtained by fitting the binding curves.

### 4.6 CRISPRi-mediated silencing of MSMEG_3124 affects *M. smegmatis* growth

To assess the functional importance of SufC ATPase in *M. smegmatis*, MSMEG_3124 was targeted using CRISPRi-mediated gene silencing, and bacterial growth was evaluated by spot-dilution assay. The MSMEG_3124-targeting CRISPRi strain exhibited reduced growth compared with the pLJR962 control upon increasing tetracycline concentrations. The growth impairment intensified at higher tetracycline concentrations, suggesting a tetracycline-dependent effect of CRISPRi-mediated repression of MSMEG_3124 (Figure 6), supporting an essential role of SufC.

**Figure 6.**
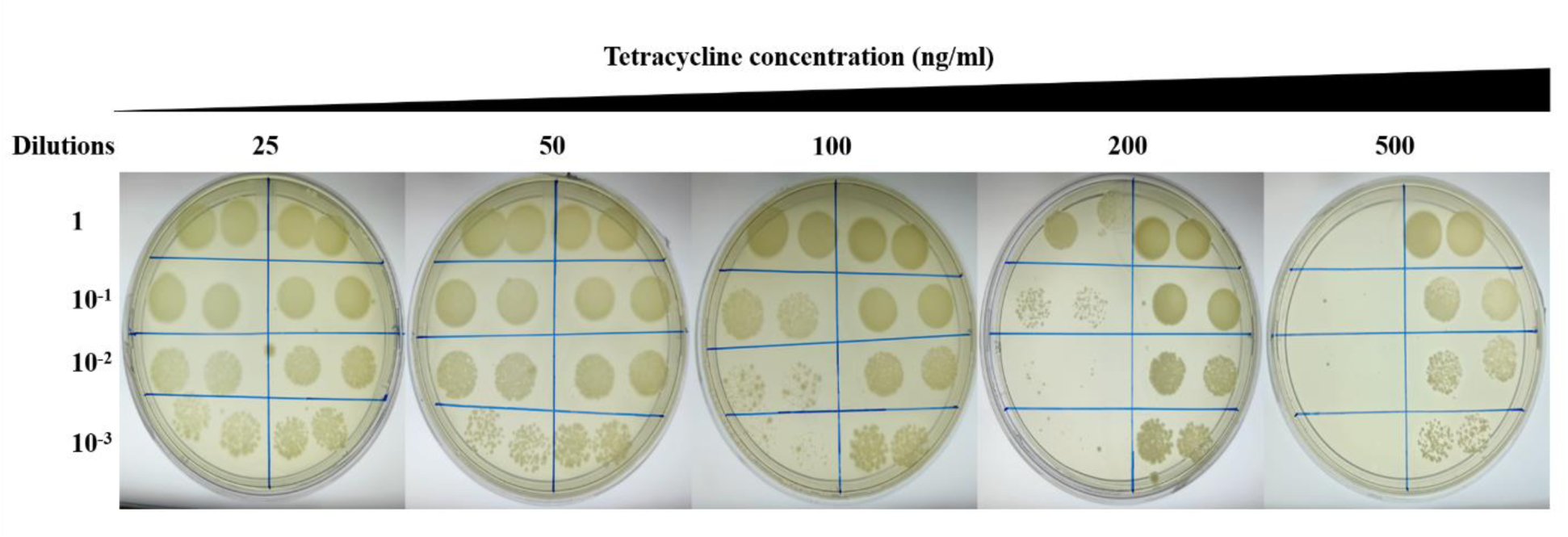
CRISPRi-mediated silencing of MSMEG_3124 in *M. smegmatis*. Growth of *M. smegmatis* carrying MSMEG_3124 targeting CRISPRi (left) and the pLJR962 vector control (right) was assessed by spot-dilution assay at increasing tetracycline concentrations (25, 50, 100, 200, and 500 ng/mL). Serial dilutions (10⁰ – 10⁻³) of the cultures were spotted onto the respective plates to evaluate the effect of MSMEG_3124 silencing on bacterial growth.

### 4.7 Generation of 3D protein structure and binding pocket analysis

The model structures of SufC from *M. tb* (Rv1463) and *M. smegmatis* (MSMEG_3124) were generated using the SWISS-MODEL server. Structural validation using MolProbity analysis showed that the Rv1463 model possessed a MolProbity score of 0.73, with 97.73% of residues located in favored Ramachandran regions. In contrast, the MSMEG_3124 structure had a MolProbity score of 0.74, with 98.04% of residues present in favored regions. These findings demonstrate the reliability of both models for further structural analysis.

Structural analysis of Rv1463 revealed a conserved ATP-binding site, which is characteristic of ABC transporter ATPases. The nucleotide-binding region is located on the Walker-A motif that spans amino acids 43-50, whereas the residues Pro-44, Asn-45, Gly-46, Ser-47, Gly-48, Lys-49, and Ser-50 form a conserved loop that interacts with the phosphate groups of ADP/ATP. The conserved Lys-49 faces the β and γ phosphates, whereas Gly-46, Ser-47, Ser-50, and Thr-51 participate in hydrogen bonding, which helps in stabilizing the nucleotide binding. Coordination with magnesium ion makes the nucleotide binding more stable (Figure 7C). However, other regions that lie outside the Walker-A region also play an essential role in stabilizing the nucleotide-binding pocket. Residue Glu-176, which corresponds to the Walker-B motif, is located distally from the phosphates and is directed for catalytic coupling rather than direct nucleotide binding, while the conserved amino acid, Gln-94, may help in stabilization of nucleotide orientation via polar interactions, even though they lie outside the classical ATPase motifs. Moreover, other peripheral residues, i.e., Val-13, Ile-25, Tyr-54, Tyr-62, and Lys-61, contribute to the formation of the ATP-binding groove by van der Waals and electrostatic interactions.

**Figure 7.**
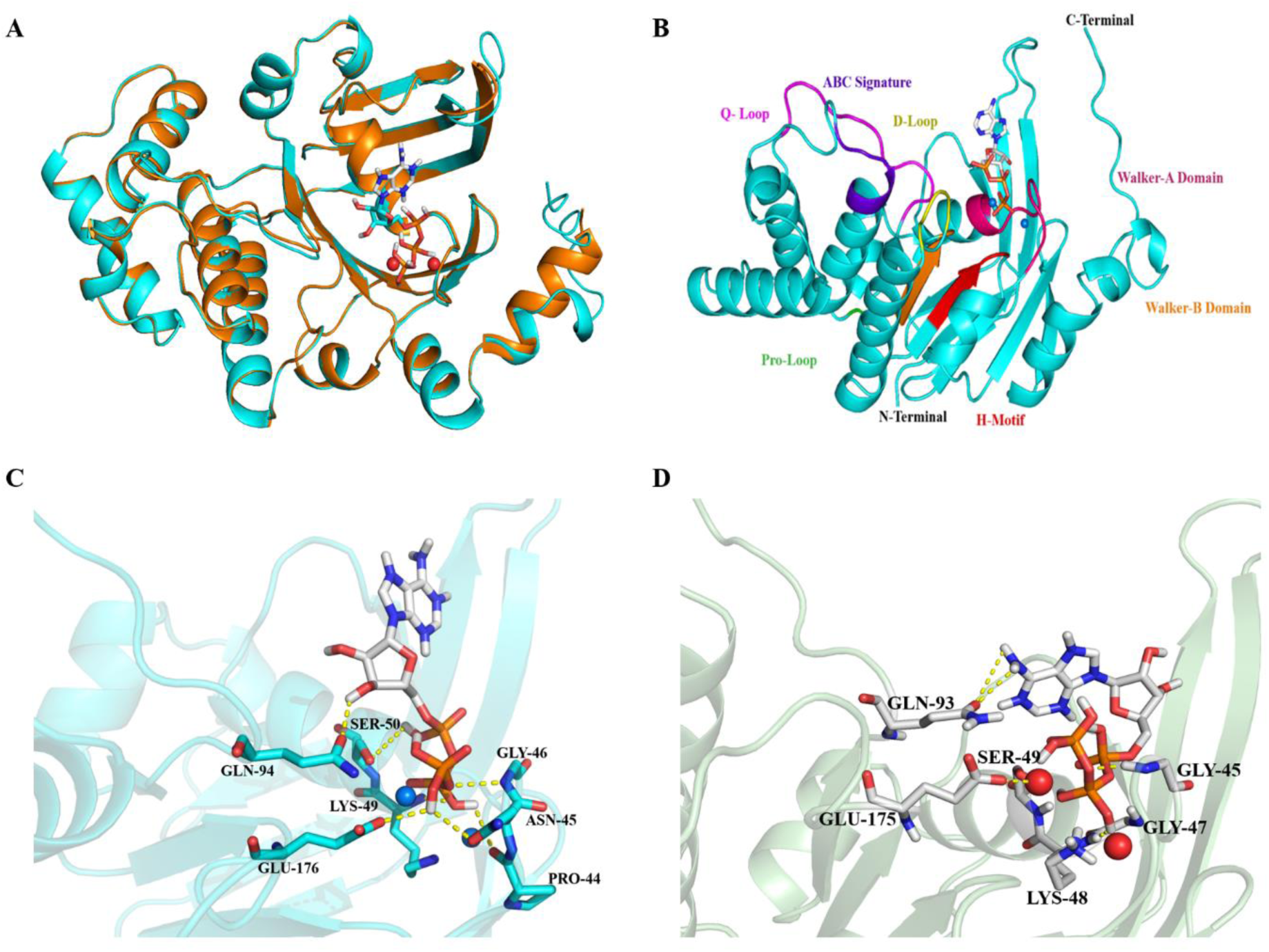
Structural characterization and ATP-binding pocket analysis of Mycobacterial SufC homologs. (A) Structural superposition of the Rv1463 (cyan) model of MSMEG_3124 (orange) model. (B) The Rv1463 model represents the conserved ATPase motifs, including the Walker-A motif, Walker-B motif, Q-loop, D-loop, H-motif, Pro-loop, and ABC signature region. (C) The three-dimensional view of ATP bound within the nucleotide-binding pocket of Rv1463, showing key interacting residues involved in ATP coordination. (D) Three-dimensional view of ATP bound within the nucleotide-binding pocket of MSMEG_3124, showing key interacting residues involved in ATP coordination.

**Figure 8.**
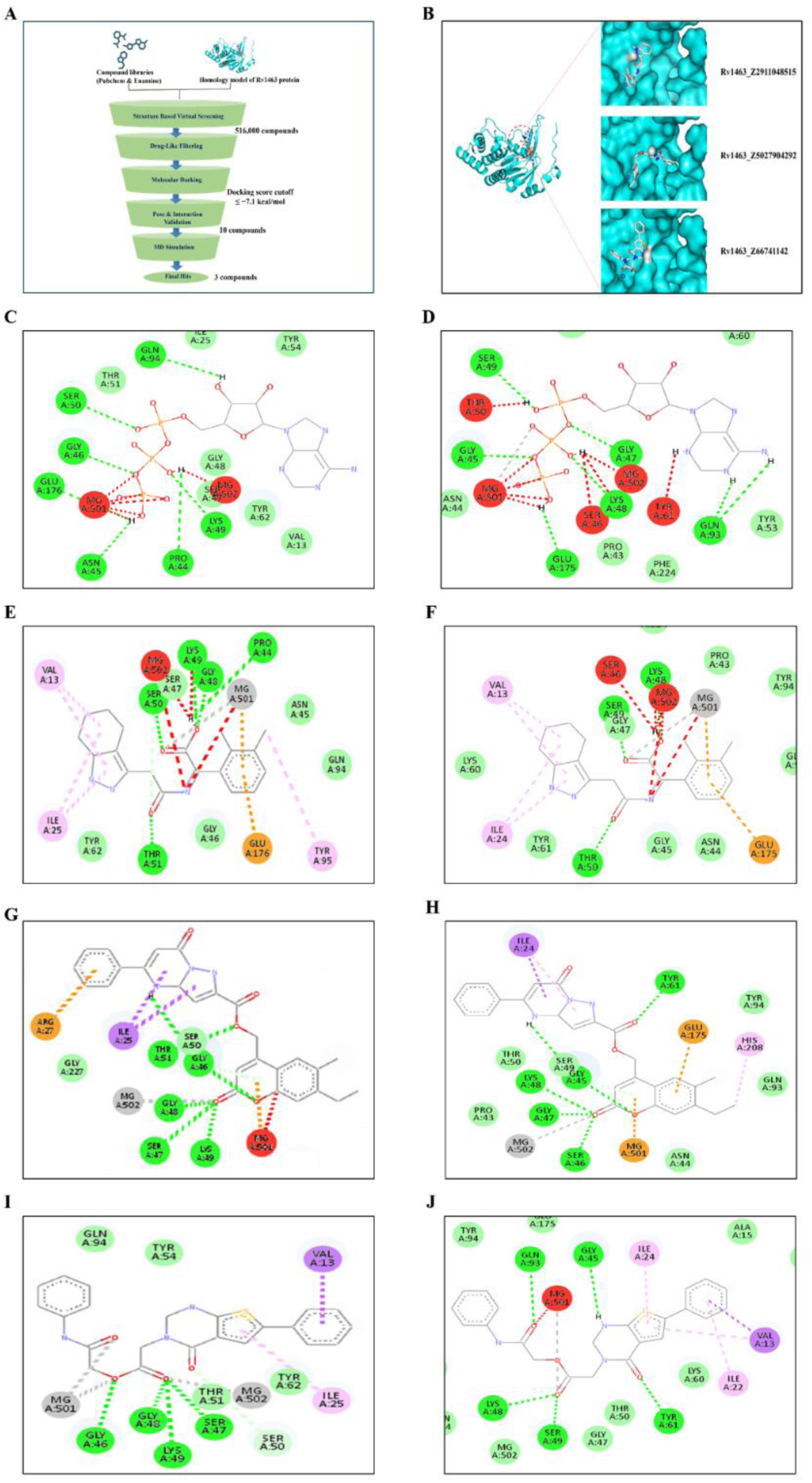
Structure-based virtual screening and comparative molecular docking analysis of mycobacterial SufC proteins. **(A)** Workflow of the structure-based virtual screening strategy used to identify potential SufC-binding compounds from the PubChem and Enamine libraries. (B) Binding poses of the three shortlisted compounds, Z2911048515, Z5027904292, and Z66741142, within the ATP-binding pocket of Rv1463. (C–D) Two-dimensional interaction maps of ATP with Rv1463 (C) and MSMEG_3124 (D). (E–F) Interaction maps of Z2911048515 with Rv1463 (E) and MSMEG_3124 (F). (G–H) Interaction maps of Z5027904292 with Rv1463 (G) and MSMEG_3124 (H). (I–J) Interaction maps of Z66741142 with Rv1463 (I) and MSMEG_3124 (J).

**Figure 9.**
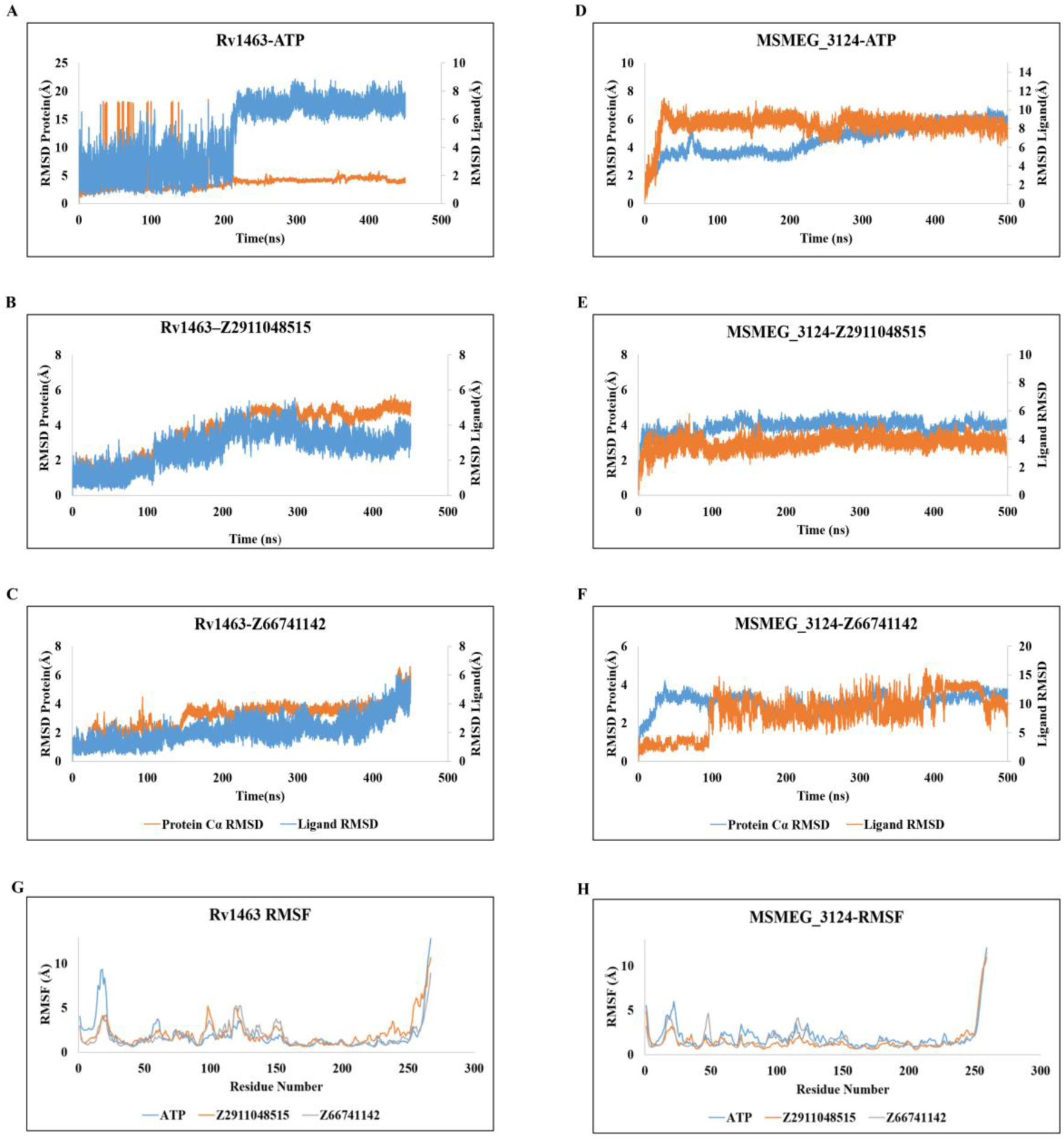
Molecular dynamics simulation analysis of Rv1463 and MSMEG_3124 complexes. (A–C) Protein Cα RMSD and ligand RMSD profiles of Rv1463 in complex with ATP (A), Z2911048515 (B), and Z66741142 (C) during 500-ns molecular dynamics simulations. (D–F) Corresponding RMSD profiles of MSMEG_3124 in complex with ATP (D), Z2911048515 (E), and Z66741142 (F). (G–H) Residue-wise RMSF profiles of Rv1463 (G) and MSMEG_3124 (H) in the presence of ATP, Z2911048515, and Z66741142.

A similar configuration of the ATP-binding site was also found for the MSMEG_3124 structure, where the geometry of the Walker-A loop, Mg²⁺ binding, and placement of supportive amino acids were preserved. The conservation of nucleotide binding pocket and the interacting residues among homologous proteins indicates a similar mechanism for nucleotide binding.

### 4.8 Molecular docking and virtual screening of SufC ATP-binding pocket

Structure-based molecular docking was performed to identify small-molecule compounds targeting the ATP-binding pocket of SufC. The docking protocol was initially validated using ATP and ADP as reference ligands to confirm accurate identification of the nucleotide-binding cleft and establish a reference interaction profile. Both ATP and ADP occupied the conserved ATP-binding region defined by the Walker-A motif (residues 43–50).

Both SufC protein models reproduced key interactions characteristic of canonical ABC ATPases, including hydrogen bonding with Ser-47, Gly-48, Lys-49, Ser-50, and Thr-51. These observations confirmed the reliability of the docking setup and the accuracy of the defined nucleotide-binding pocket. Following validation, approximately 5 × 10⁵ drug-like molecules from the Enamine and PubChem databases were subjected to virtual screening using AutoDock Vina. Virtual screening identified multiple compounds with predicted binding energies ranging from −7.5 to −8.7 kcal/mol, comparable to or exceeding those obtained for ATP and ADP. Based on their docking scores, the top-ranked 50 compounds were selected for further evaluation. Independent docking runs using different random seeds and exhaustiveness values confirmed reproducible binding orientations within the ATP-binding pocket. Further compounds were evaluated for pose reproducibility, interaction profiles, PAINS characteristics, and ligand efficiency, resulting in 10 compounds as the most promising candidates for downstream analysis.

The interaction analysis of the shortlisted compounds showed that the identified top-10 compounds occupied the ATP-binding pocket and established hydrogen-bonding networks with residues of the Walker-A motif. Additional stabilizing interactions involving residues such as Pro-44, Gly-46, and Tyr-62 further contributed to ligand accommodation within the catalytic pocket. A subset of ligands also displayed Mg²⁺-associated coordination patterns. Subsequent filtering based on docking score, interaction consistency, pose reproducibility, PAINS filtering, and ligand efficiency yielded a refined set of compounds with stable occupancy and favorable physicochemical properties. Among these, Z2911048515, Z66741142, and Z5027904292 demonstrated the most favorable interaction profiles and were selected for downstream analysis.

To further assess conservation of ligand recognition across mycobacterial homologs, the shortlisted top-10 compounds were additionally docked against MSMEG_3124 model. The compounds displayed comparable binding orientations and conserved interaction patterns within the ATP-binding pocket of MSMEG_3124, supporting structural conservation of the nucleotide-binding environment across pathogenic and non-pathogenic mycobacterial SufC homologs. These selected complexes were subsequently subjected to molecular dynamics simulations to evaluate binding stability and the persistence of interactions under dynamic conditions.

### 4.9 Selectivity analysis against Human ABC ATPases

To evaluate selectivity, three shortlisted compounds (CID: Z2911048515, Z66741142, and Z5027904292) were further docked against a representative human ABC ATPase (PDB ID: 3ZDQ). Structural models of the nucleotide-binding domains were prepared and subjected to the same docking protocol used for Rv1463 and MSMEG_3124 to ensure consistency in comparative analysis. In contrast to their stable binding within the SufC ATP-binding pocket, the selected compounds exhibited reduced binding affinity and inconsistent docking poses for 3ZDQ. Moreover, the compounds failed to maintain stable positioning within the nucleotide-binding pocket and did not reproduce key interactions typically associated with ATP binding. Notably, interactions with critical residues of the Walker-A motif, which are essential for nucleotide coordination, were either weak or absent. Furthermore, pose reproducibility analysis revealed variability across docking runs, indicating a lack of stable binding modes in the 3ZDQ structure. In several cases, the compounds were partially displaced from the canonical binding pocket, suggesting poor accommodation within 3ZDQ active site. These observations highlight a clear distinction in binding behavior between human ABC ATPase and mycobacterial SufC ATPases. Despite the conserved nature of the ATP-binding motifs, differences in the surrounding pocket environment and residue composition appear to contribute to selective ligand recognition. To further validate these findings, the protein-ligand complexes of the selected compounds with both SufC ATPase and a human ATPase were subjected to molecular dynamics simulations to assess binding stability and the persistence of interaction under dynamic conditions.

### 4.10 Molecular dynamics simulation results

#### Short molecular dynamics simulations

To evaluate the stability of the docked complexes, 50 ns molecular dynamics (MD) simulations were performed for ATP-bound Rv1463 and the ligand-bound complexes identified through virtual screening. RMSD and RMSF analyses demonstrated variable stabilization profiles among the screened compounds. Several ligands showed increased fluctuations and weak retention within the ATP-binding pocket. In contrast, complexes containing Z2911048515, Z66741142, and Z5027904292 displayed comparatively stable RMSD trajectories and persistent catalytic-site occupancy throughout the simulation period. Residue-level fluctuations were primarily restricted to loop and terminal regions, while residues constituting the ATP-binding pocket remained comparatively stable. Based on their favorable interaction profiles and structural stability, Z2911048515, Z66741142, and Z5027904292 were selected for extended 500 ns simulations.

#### Long-term molecular dynamics simulations of Rv1463 complexes

Extended 500 ns simulations were performed for ATP-bound Rv1463 and the three selected ligand-bound complexes to evaluate long-term conformational stability and interaction persistence. The ATP-bound control system maintained stable catalytic-site interactions throughout the trajectory, including persistent Mg²⁺ coordination and conserved contacts involving Gly-46, Ser-47, Ser-50, Gln-94, Asp-175 and Glu-176. Secondary structure analysis further confirmed preservation of the overall protein fold during simulation.

Among the ligand-bound systems, the Rv1463-Z2911048515 complex exhibited the most stable interaction profile, characterized by sustained Mg²⁺ coordination, persistent hydrogen-bond and ionic interactions, and stable occupancy within the ATP-binding pocket. The Z66741142-bound complex showed intermediate stability with comparatively higher ligand mobility during later stages of the trajectory. In contrast, the Z5027904292-bound complex displayed reduced interaction persistence and increased solvent-mediated contacts, suggesting comparatively weaker stabilization within the catalytic pocket. Comparative analyses of RMSF, SASA, and radius of gyration demonstrated that all complexes maintained overall structural integrity throughout the simulations.

#### Comparative analysis with *M. smegmatis* SufC

Additional 500 ns simulations were performed using the homologous MSMEG_3124 ATPase system in complex with ATP, Z2911048515, and Z66741142 to evaluate the conservation of interactions within a non-pathogenic mycobacterial homolog. The ATP-bound MSMEG_3124 complex displayed stable catalytic-site occupancy and conserved Mg²⁺-associated interactions throughout the trajectory.

Both Z2911048515 and Z66741142 maintained interactions within the ATP-binding pocket of MSMEG_3124, although their persistence was comparatively reduced compared with the Rv1463 complexes. The MSMEG_3124-Z2911048515 complex showed improved interaction stability and more sustained catalytic-site contacts compared with the MSMEG_3124-Z66741142 complex, which exhibited greater ligand mobility and weaker interaction persistence during later stages of the trajectory. Overall, the simulations demonstrated conservation of major catalytic interaction patterns across pathogenic and non-pathogenic mycobacterial SufC homologs while highlighting differences in ligand stabilization dynamics.

#### Comparative analysis with homologous human ATPase

To investigate ligand selectivity, 500 ns simulations were performed using the homologous human ATPase structure (3ZDQ) in complex with ATP, Z2911048515, and Z66741142. Compared with the Rv1463 protein, the human ATPase complexes exhibited greater conformational flexibility, reduced persistence of Mg²⁺-associated interactions, and weaker ligand stabilization within the nucleotide-binding pocket. Although ATP maintained catalytic-site association throughout the simulation, ligand-bound complexes exhibited greater mobility and less stable interaction networks than their mycobacterial counterparts. Among the tested ligands, Z2911048515 retained comparatively stronger interactions than Z66741142 within the human ATPase environment.

## 5. Discussion

The SUF pathway is vital for iron-sulfur (Fe-S) cluster biogenesis in *M. tb*, as it lacks the ISC and NIF systems. The SufC proteins from Rv1463 in *M. tb* and MSMEG_3124 in *M. smegmatis* are ATPase components of the SUF system, which are involved in ATP-binding and hydrolysis, and function within the SufB₂CD scaffold ^22,24,26^. The CRISPR interference experiments demonstrated that growth impairment became more pronounced with increasing tetracycline concentrations, implicating the repression of MSMEG_3124 (Figure 6) and reinforcing the essential function of SufC, consistent with findings from previous studies. SufC plays a vital role in the maintenance of iron-sulfur (Fe-S) cluster homeostasis, particularly under stressors such as oxidative and nitrosative stress that disrupt cluster integrity ^15,16,46^. Despite its essential role in bacterial viability and fitness, mycobacterial SufC proteins remain poorly characterized.

In this study, SufC structural models were prepared for Rv1463 and MSMEG_3124. These models maintain the conserved nucleotide-binding architecture characteristic of bacterial SufC proteins and other ABC-type ATPases ^29,30^. Through structure-guided virtual screening at the putative ATP-binding site, three compounds from the Enamine library, i.e., Z2911048515, Z66741142, and Z5027904292, showed favorable predicted binding scores. The molecular dynamics simulations indicated that the compounds could maintain stable interactions within the predicted binding pocket, although the interaction patterns and stability profiles varied across complexes.

To confirm the computational predictions, Rv1463 and MSMEG_3124 were successfully cloned and purified to over 90% homogeneity. SDS-PAGE analysis indicated that both proteins have a molecular weight of approximately 30 kDa. Size-exclusion chromatography profiles indicated that both proteins predominantly exist as monomers, aligning with previous observations for *T. thermophilus* SufC and *E. coli* SufC ^27,30^. Importantly, incubating Rv1463 and MSMEG_3124 with ATP and Mg²⁺ did not elicit a notable change in their SEC elution profiles, suggesting that ATP and Mg²⁺ do not alter the oligomeric state of SufC proteins (Supplementary Figure 3). These findings are consistent with earlier studies on *E. coli* SufC oligomerization, indicating that ATP and Mg^2+^ do not influence oligomer formation. In contrast, SufC interacts individually with SufB and SufD to facilitate the formation of the SufBC_2_D scaffold ^24,27,29^. Interestingly, the *SufC* protein in *T. martima and E. coli* is a monomer with low intrinsic ATPase activity, whereas the presence of SufB and SufD significantly increases this activity through ATP-dependent conformational changes within the SufBC₂D complex ^27,31,47^.

To evaluate differences in secondary-structure composition between the two proteins, a comparison of Far-UV CD spectra reveals that MSMEG_3124 has a higher α-helical content than Rv1463. Moreover, the addition of ATP to the protein solution led to minor changes in the CD spectra of both proteins, indicating that nucleotide interaction induces minor alterations in secondary structure (Figure 10). The CD spectra obtained after incubation of Z2911048515 or Z66741142 with their respective mycobacterial homologs displayed unique spectral profiles. Incubation of Rv1463 with Z2911048515 or Z66741142 induced minor changes in the far-UV CD spectrum. In contrast, incubation of MSMEG_3124 with Z2911048515 or Z66741142 exhibited pronounced spectral alterations, characterized by a significant decrease in α-helical content, suggesting that the compounds preferentially impact the secondary structure of MSMEG_3124 compared to Rv1463. Furthermore, both proteins contain a single tryptophan residue, so fluorescence spectroscopy was used to investigate the impact of ATP and Enamine compounds (Z2911048515 or Z66741142). Surprisingly, incubation with Z66741142 resulted in a notable reduction in tryptophan fluorescence intensity in both proteins, indicating that the compound induced conformational changes that reduced tryptophan emission. Both CD and fluorescence spectroscopy results indicate that the selected compounds altered secondary-structure content or structural conformation. Surprisingly, these structural changes had only modest effects on ATPase activity, indicating that a ligand-induced alteration of secondary structure does not translate into efficient inhibition of ATP hydrolysis. However, the Z2911048515 compound showed slight inhibition, whereas Z66741142 showed negligible inhibition, possibly due to its solubility in the reaction buffer. However, these compounds may contribute as preliminary scaffolds and could be further improved to increase their inhibitory efficacy.

**Figure 10.**
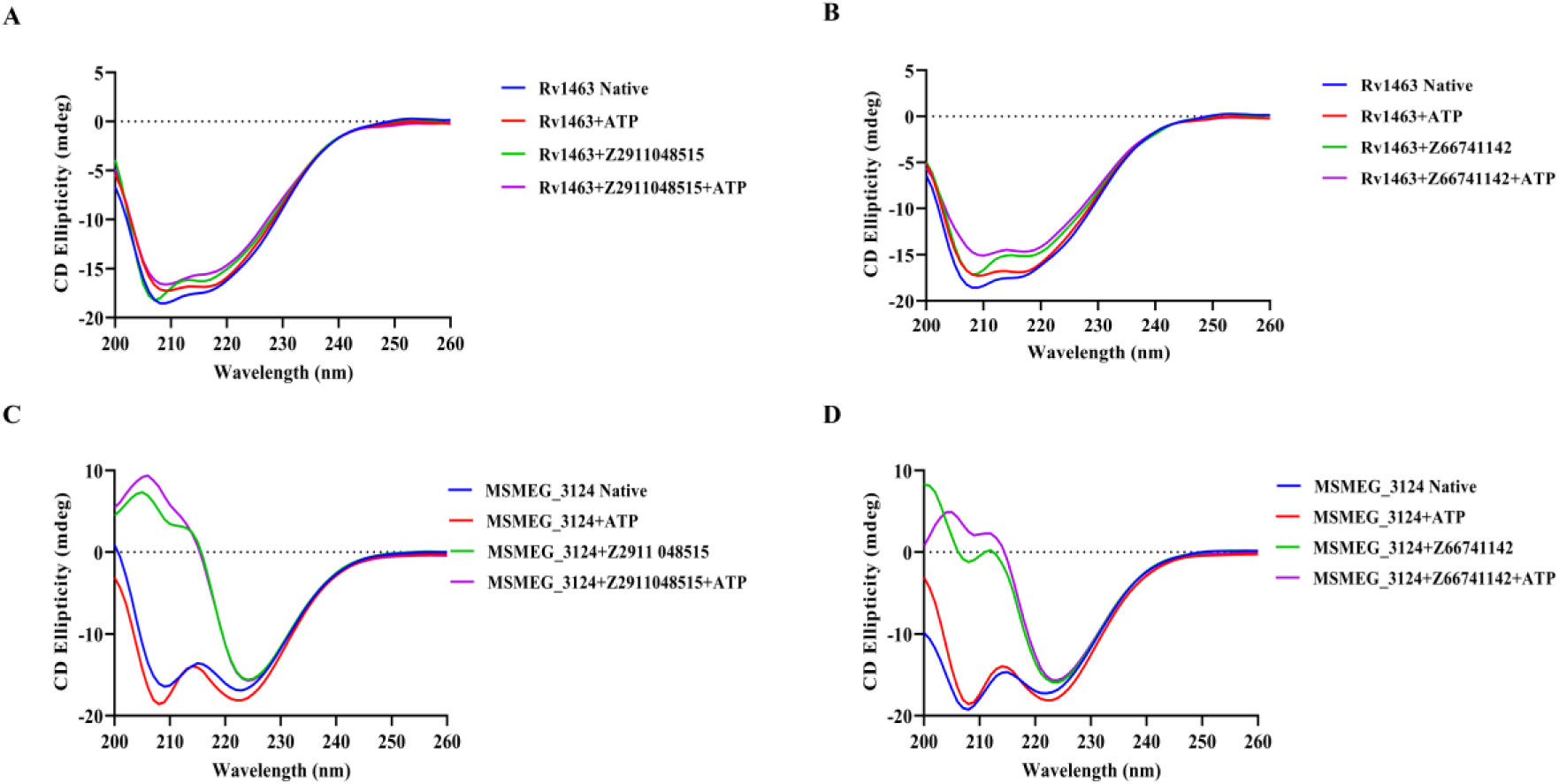
Ligand-induced secondary structure changes in mycobacterial SufC proteins analyzed by circular dichroism spectroscopy. **(A)** Far-UV CD spectra of recombinant *Mycobacterium tuberculosis* SufC (Rv1463) in the native state and in the presence of ATP, Z2911048515, and Z2911048515 in the presence of ATP. **(B)** Far-UV CD spectra of Rv1463 in the native state and in the presence of ATP, Z66741142, and Z66741142 in the presence of ATP. **(C)** Far-UV CD spectra of recombinant *Mycolicibacterium smegmatis* SufC (MSMEG_3124) in the native state and in the presence of ATP, Z2911048515, and Z2911048515 in the presence of ATP. **(D)** Far-UV CD spectra of MSMEG_3124 in the native state and in the presence of ATP, Z66741142, and Z66741142 in the presence of ATP.

**Figure 11.**
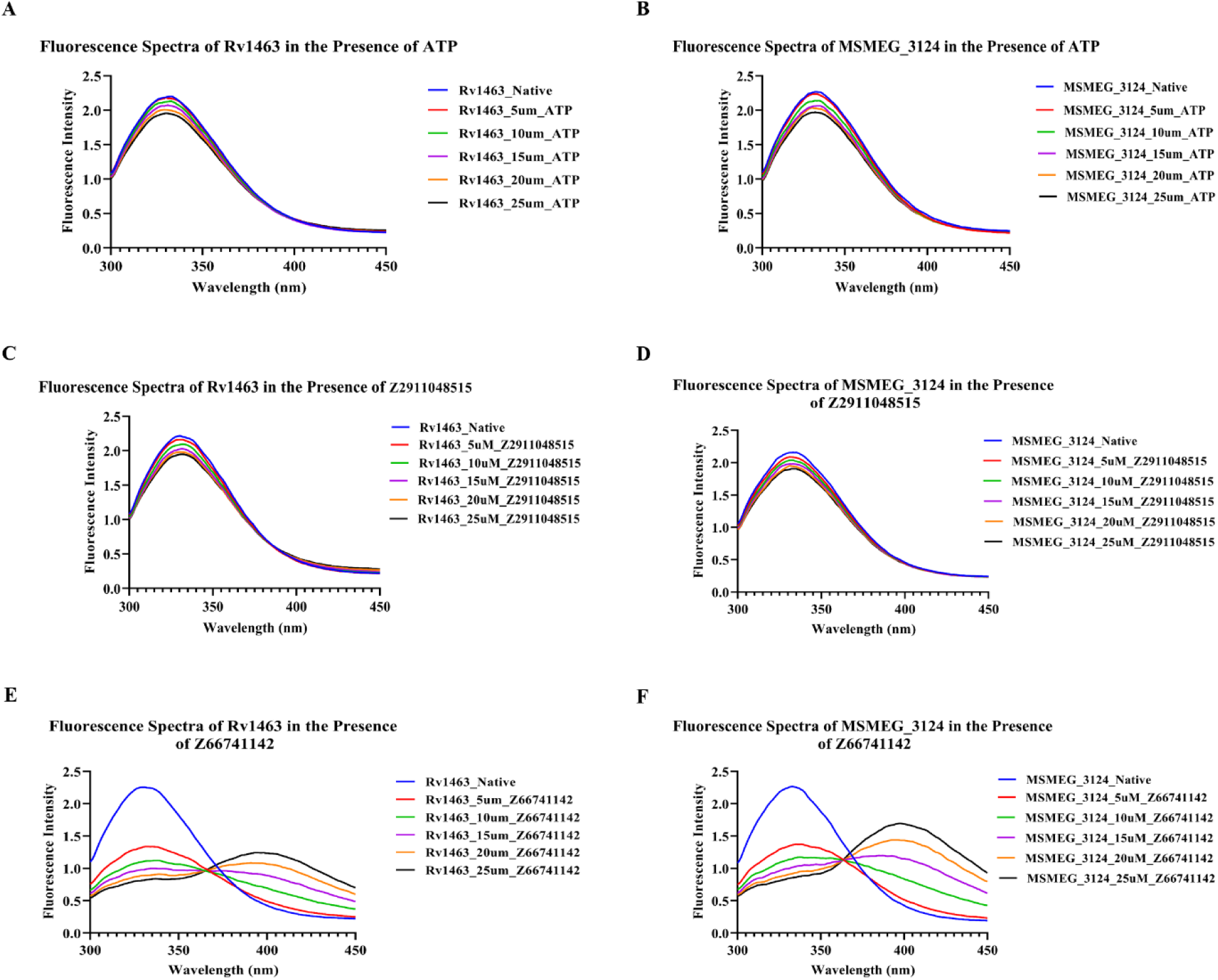
Intrinsic fluorescence analysis of ligand-induced changes in mycobacterial SufC proteins. (A–B) Fluorescence spectra of Rv1463 (A) and MSMEG_3124 (B) in the presence of increasing concentrations of ATP. (C–D) Fluorescence spectra of Rv1463 (C) and MSMEG_3124 (D) in the presence of increasing concentrations of Z2911048515. (E–F) Fluorescence spectra of Rv1463 (E) and MSMEG_3124 (F) in the presence of increasing concentrations of Z66741142.

Although the overall structures of Rv1463 and MSMEG_3124 are closely related, they differ in their kinetic properties. Rv1463 exhibited a lower Km for ATP, whereas MSMEG_3124 displayed a higher Vmax and Kcat values. This could be beneficial for mycobacteria as the lower Km of Rv1463 might facilitate more efficient ATP utilization at lower ATP concentrations, which may be particularly beneficial for the Fe–S cluster biogenesis when energy resources are limited. The observed kinetic divergences among SufC homologs may be attributed to slight differences in their ATP-binding and catalytic domains, which are likely modulated by local structural dynamics and help *M. tb* adapt to unfavorable conditions, including iron deprivation and oxidative stress, which disrupts iron-sulfur (Fe–S) cluster homeostasis.

In summary, these findings provide a comparative biochemical characterization of mycobacterial SufC homologs and lay the foundation for optimizing compounds that target SufC and ultimately elucidating their role in mycobacterial physiology.

## 7. Acknowledgments

The authors would like to express their gratitude to Professor William R. Jacobs from the Department of Microbiology and Immunology and the Howard Hughes Medical Institute at Albert Einstein College of Medicine, Bronx, New York, USA, for providing the pYUB1062 expression vector, and Dr. Prem Chandra Kaushal, RCB, Gurugram, India, for providing *M. smegmatis* mc^2^155 expression strain. The plasmid Rv1463-pANT7-cGST (clone MtCD00815348) was obtained from the DNASU Plasmid Repository (Arizona State University, Tempe, AZ, USA). The authors are also grateful to the Department of Life Sciences, Shiv Nadar Institution of Eminence (SNIoE), for providing access to the Central Instrumentation Facility (CIF). The authors are also thankful to the Department of Life Sciences, SNIoE for financial support to Rajan Vyas and for a Ph.D. fellowship to Anu Ghodpage.

## 8. Author Contributions

**AG.:** Investigation, Methodology, Data Curation, Formal Analysis, and Writing – Original Draft, **AD.:** MD simulation, Review and editing, **AS.:** Supervision of MD simulation, Review & Editing the manuscript, **R.V.:** Supervision, Methodology, Project Administration, Funding Acquisition, and Writing – Review & Editing.

