## Supplementary information for "Comparative Biochemical and Biophysical Insights into SufC ATPases from *Mycobacterium tuberculosis* and *Mycolicibacterium smegmatis mc^2^155*"

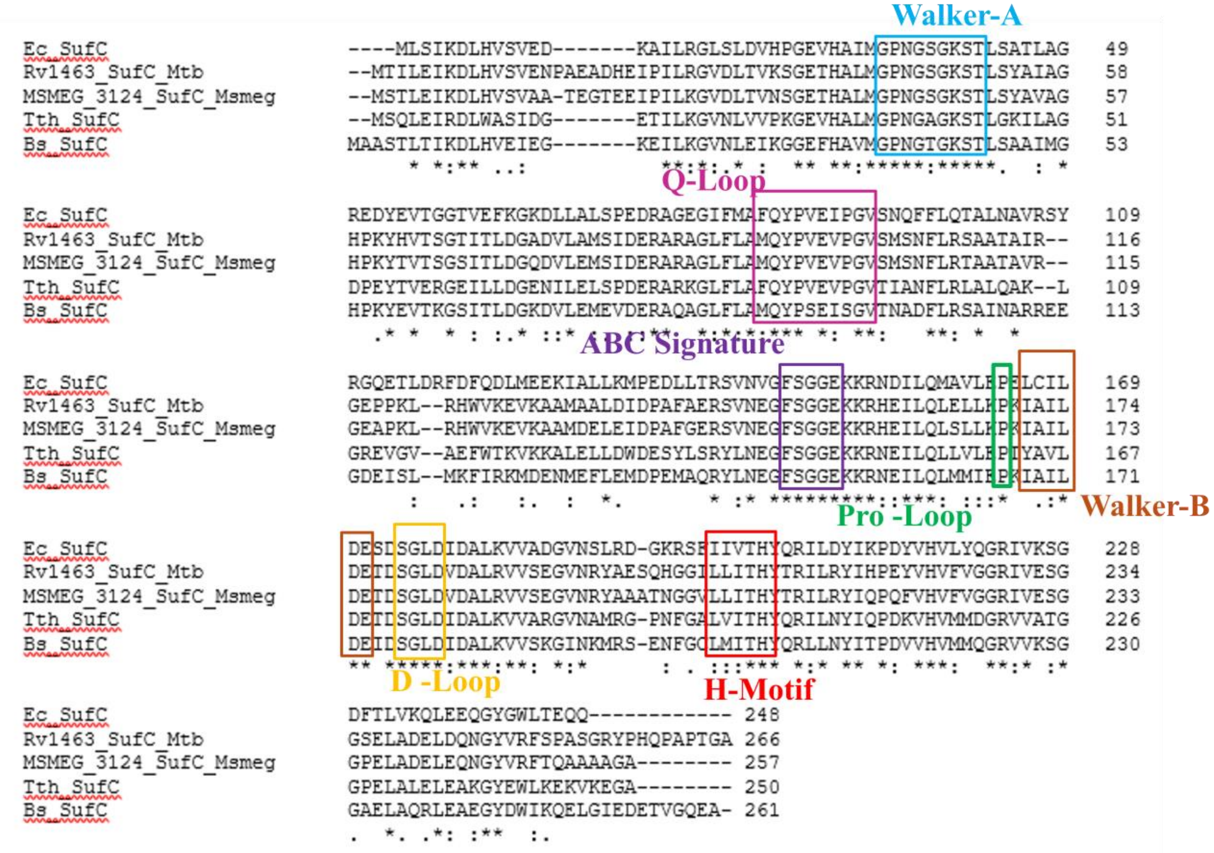


**Supplementary Figure 1.** Multiple sequence alignment of SufC homologs. Highlighted regions indicate conserved ABC ATPase motifs (Walker-A, Walker-B, Q-loop, H-loop, and signature-like loop).

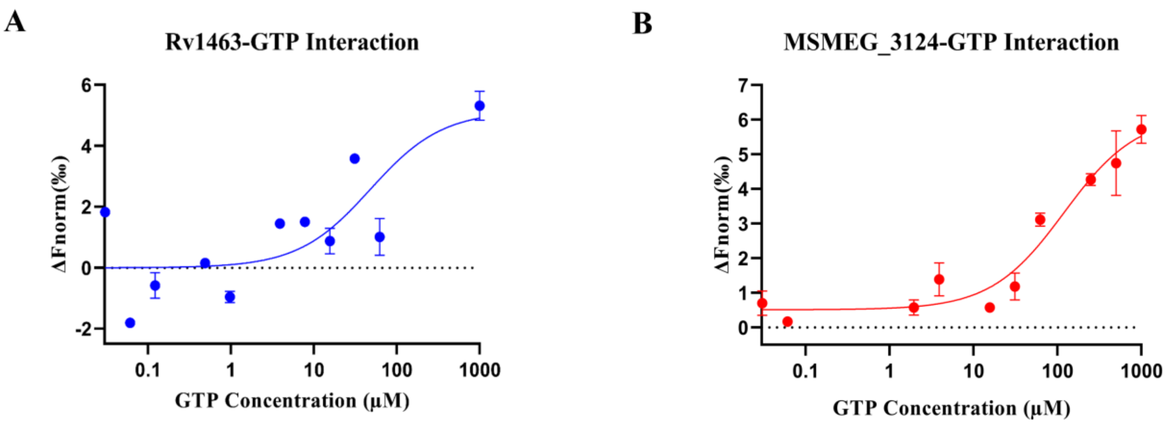


**Supplementary Figure 2. Nucleotide binding by SufC homologs determined by microscale thermophoresis (MST). (A)** Binding of ATP and GTP to Rv1463. **(B)** Binding of ATP and GTP to MSMEG_3124. The change in normalized fluorescence (ΔFnorm) was plotted as a function of nucleotide concentration, and Kd values were obtained by fitting the binding curves.

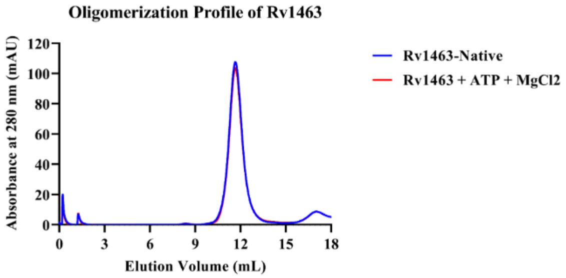


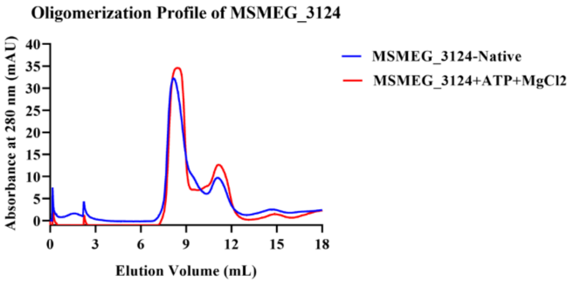


**Supplementary Figure 3. Analytical size-exclusion chromatography profiles of mycobacterial SufC proteins in the presence of ATP. (A)** Elution profiles of Rv1463 in the native state and in the presence of ATP and $MgCl_2$. **(B)** Elution profiles of MSMEG_3124 in the native state and in the presence of ATP and $MgCl_2$. Absorbance was monitored at 280 nm.

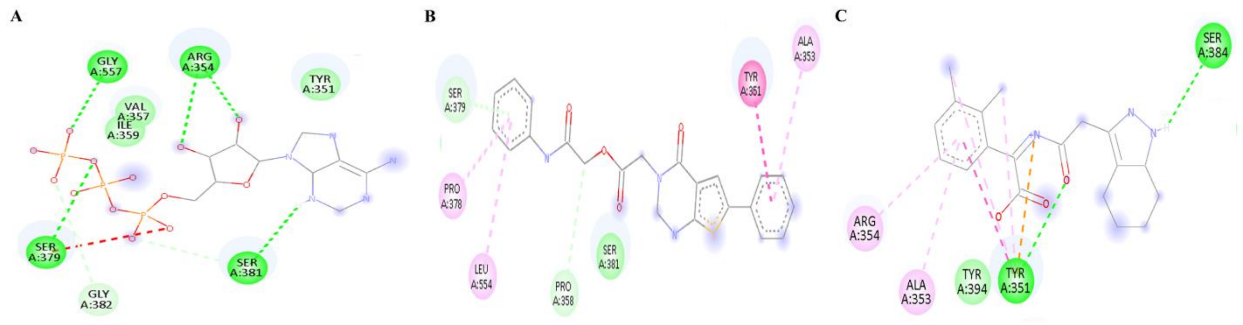


**Supplementary Figure 4. Two-dimensional interaction maps of ATP and selected compounds docked to 3ZDQ. (A)** ATP, **(B)** Z66741142, and **(C)** Z2911048515 bound at the ATP-binding site of 3ZDQ. The interaction maps were generated using Discovery Studio Visualizer.

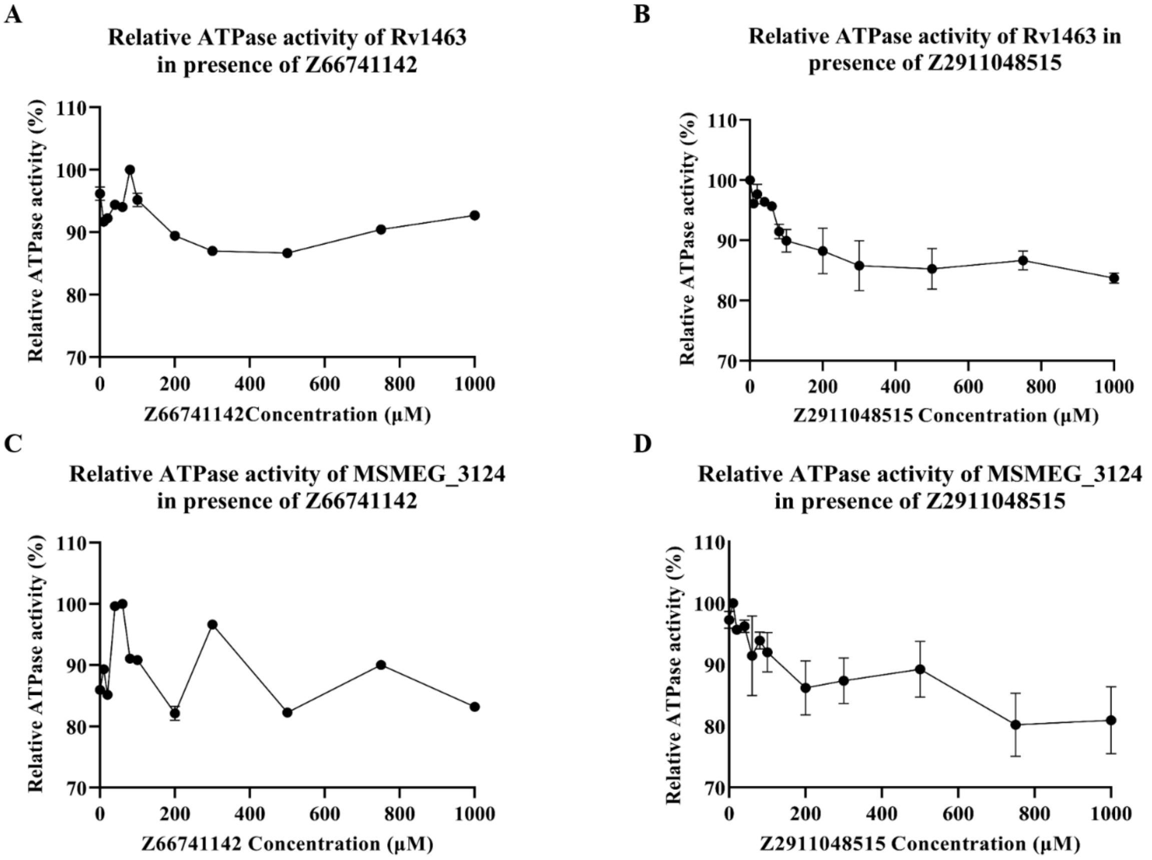


**Supplementary Figure 5. Relative ATPase activity of mycobacterial SufC proteins in the presence of selected compounds.** (A) Relative ATPase activity of recombinant *Mycobacterium tuberculosis* SufC (Rv1463) in the presence of increasing concentrations of Z66741142. (B) Relative ATPase activity of Rv1463 in the presence of increasing concentrations of Z2911048515. (C) Relative ATPase activity of recombinant *Mycolicibacterium smegmatis* SufC (MSMEG_3124) in the presence of increasing concentrations of Z66741142. (D) Relative ATPase activity of MSMEG_3124 in the presence of increasing concentrations of Z2911048515. Relative ATPase activity is expressed as a percentage of untreated control. Data represent the mean ± SD from three independent experiments.

### Supplementary Table S1. Primers used in this study

| Primers | Sequence |
| --- | --- |
| Rv1463_pYUB_F1 | 5’-GGATATATACATATGACCATTTTGGAAATTAAGGACC-3’ |
| Rv1463_pYUB_R1 | 5’-TATAAGTATAAGCTTTTAGTGATGGTGATGGTGATGGGCTCCGGTTGGCGCGGGT TGG-3’ |
| MSMEG_3124_F1 | 5’- GGATATATACATATGAGCACCCTGGAAATCAAGGACC-3’ |
| MSMEG_3124_R1 | 5’-TATAAGTATAAGCTTTTAGTGATGGTGATGGTGATGGGCCCCTGCGGCTGCTGC CTGCG-3’ |

**<u>Supplementary Table S2.</u> sgRNAS used in CRISPRi study**

| sgRNA | Sequence | Fold Repression |
|---|---|---|
| MSMEG_3124_N1_T | 5’-AAACGGCGGTGAGAAGAAGCGCCACG-3’ | 216.7 |
| MSMEG_3124_N1_B | 5’-GGGACGTGGCGCTTCTTCTCACCGCC-3’ | 216.7 |
| MSMEG_3124_N2_T | 5’-GGGAGCACCGCGGTCGCGGCCGTGCG-3’ | 145.2 |
| MSMEG_3124_N2_B | 5’-AAACCGCACGGCCGCGACCGCGGTGC-3’ | 145.2 |
| MSMEG_3124_N3_T | 5’-GGGATTGGGCTTGAGCAACGACAGCT-3’ | 82.2 |
| MSMEG_3124_N3_B | 5’-AAACAGCTGTCGTTGCTCAAGCCCAA-3’ | 82.2 |
